# Efficacy of SHP2 phosphatase inhibition in cancers with nucleotide-cycling oncogenic RAS, RAS-GTP dependent oncogenic BRAF and NF1 loss

**DOI:** 10.1101/188730

**Authors:** Robert J. Nichols, Franziska Haderk, Carlos Stahlhut, Christopher J. Schulze, Golzar Hemmati, David Wildes, Christos Tzitzilonis, Kasia Mordec, Abby Marquez, Jason Romero, Daphne Hsieh, Gert Kiss, Elena S. Koltun, Adrian L. Gill, Mallika Singh, Mark A. Goldsmith, Jacqueline A. M. Smith, Trever G. Bivona

## Abstract

Oncogenic alterations in the RAS-RAF-MEK-ERK pathway, including mutant forms of KRAS, BRAF, and loss of the tumor suppressor and RAS GTPase-activating protein (GAP) NF1, drive the growth of a wide spectrum of human cancers. While BRAF and MEK inhibitors are effective in many patients with oncogenic BRAF V600E, there are no effective targeted therapies for individuals with cancers driven by other pathway alterations, including oncogenic KRAS, non-V600E BRAF, and NF1 loss. Here, we show that targeting the PTPN11/SHP2 phosphatase with a novel small molecule allosteric inhibitor is effective against cancers bearing nucleotide-cycling oncogenic RAS (e.g. KRAS G12C), RAS-GTP dependent oncogenic BRAF (e.g. class 3 BRAF mutants), or NF1 loss in multiple preclinical models in vitro and in vivo. SHP2 inhibition suppressed the levels of RAS-GTP and phosphorylated ERK in these models and induced growth inhibition. Expression of a constitutively active mutant of the RAS guanine nucleotide exchange factor (GEF) SOS1 rescued cells from the effects of SHP2 inhibition, suggesting that SHP2 blockade decreases oncogenic RAS-RAF-MEK-ERK signaling by disrupting SOS1-mediated RAS-GTP loading. Our findings illuminate a critical function for SHP2 in promoting oncogenic RAS activation and downstream signaling in cancers with nucleotide-cycling oncogenic RAS, RAS-GTP dependent oncogenic BRAF, and NF1 loss. SHP2 inhibition thus represents a rational, biomarker-driven therapeutic strategy to be tested in patients with cancers of diverse origins bearing these oncogenic drivers and for which current treatments are largely ineffective.

## INTRODUCTION

RAS proteins are small GTPases that operate as molecular switches. When GTP-bound, RAS can engage downstream effector proteins, including RAF, to activate the MEK-ERK (MAPK) pathway and promote cellular proliferation^1^. Activation of receptor tyrosine kinases (RTKs) leads to activation of RAS guanine nucleotide exchange factors (GEFs), such as son of sevenless homolog 1 (SOS1), to promote GTP loading of RAS and signaling, while RAS GTPase-activating proteins (GAPs) stimulate GTP hydrolysis and terminate RAS activation and signaling^2^. This homeostatic GEF/GAP cycle is disrupted by oncogenic mutations in RAS and upstream components of the pathway, and can also be bypassed by mutations in downstream components.

Oncogenic activation of the RAS/MAPK signaling pathway occurs frequently in a wide spectrum of tumor types^3^. In lung adenocarcinoma, oncogenic pathway activation most commonly arises via mutations in KRAS, but also can result from mutations in upstream or downstream components of the pathway such as RTKs or BRAF^4^. Inactivation of the tumor suppressor gene neurofibromin 1 (NF1), which encodes a RAS GAP, is an additional mechanism driving dysregulated RAS signaling in multiple tumor types^5,6^. Despite extensive clinical evaluation of RAF and MEK inhibitors across multiple tumor histologies harboring oncogenic mutations in RAS or RAF^7^, only tumors driven by BRAF^V600^ mutations have shown consistent clinical responses^8^. Identifying new therapeutic strategies to block RAS/MAPK pathway signaling and growth in tumors with oncogenic activation of the pathway is of paramount importance.

A commonly accepted paradigm is that driver mutations in the RAS/MAPK pathway are constitutively active and function independently of normal upstream regulation. A growing body of evidence suggests this may not be true in all cases. In particular, oncogenic mutations of KRAS were generally considered to be “constitutively active,” i.e. locked in a GTP-bound state. However, recent studies have demonstrated that some oncogenic mutations of KRAS, like G12C, undergo active cycling between GTP‐ and GDP-bound states and therefore remain sensitive to modulation by upstream factors^9-11^. In addition, the recently described class 3 mutations of BRAF promote RAF hetero-dimerization and signaling in a RAS-GTP dependent manner and thus also require upstream signals to drive aberrant cell growth^12,13^. Oncogenic variants of this type appear to amplify physiologic growth signals rather than drive signaling autonomously. The therapeutic strategy of limiting GTP loading of RAS in tumors with such oncogenic RAS/MAPK pathway mutations is a rational approach that has not been fully explored to date.

SHP2 (PTPN11) is a non-receptor protein tyrosine phosphatase and scaffold protein that functions downstream of multiple RTKs, integrating growth factor signals to promote RAS activation^14^. Accordingly, autosomal dominant activating mutations in PTPN11 fuel pathogenic RAS/MAPK signaling and are potent drivers of RASopathies and human cancers^15,16^. Genetic knockdown or pharmacological inhibition of SHP2 suppresses RAS/MAPK signaling and has been shown to limit the proliferation of cancer cells dependent upon a range of activated RTKs^17^, pointing to a therapeutic opportunity for SHP2 inhibition in RTK-driven cancers.

Here, we describe the use of a novel, potent, and selective SHP2 allosteric inhibitor, RMC-4550. RMC-4550 was used to explore whether inhibition of SHP2 can suppress RAS activation and proliferation in cancers that, distinct from those dependent on oncogenic RTKs, are driven by downstream mutations dependent on RAS nucleotide cycling. As a convergent node for growth factor/RTK signaling, SHP2 may be a well-positioned target for disruption of the upstream signaling events that are necessary for pathogenic RAS/MAPK signaling in a genetically-defined subset of tumors.

## RESULTS

### Identification of KRAS Mutations that Confer Sensitivity to SHP2 Inhibition

To test the hypothesis that SHP2 inhibition suppresses oncogenic RAS signaling, we sought to identify a SHP2 small molecule inhibitor. A potent, selective, allosteric inhibitor of SHP2, RMC-4550, was identified through a biochemical and cellular screen of novel compounds and subsequent structure-guided medicinal chemistry campaign to optimize potency, selectivity, and drug-like properties (manuscript in preparation). RMC-4550 inhibited the activity of full-length SHP2 enzyme with a potency of 0.58 nM, and had no activity against the SHP2 catalytic domain up to a concentration of 10 μM (Fig. S1A). RMC-4550 was highly selective for full-length SHP2 across panels comprising 15 other phosphatases, 468 kinases, and 44 cellular targets (Tables S1-2, Fig. S2).

To determine whether specific driver mutations in KRAS proteins are sensitive to SHP2-dependent regulation of RAS-GTP levels, we screened a panel of thirty-three KRAS-mutant cancer cell lines for sensitivity to RMC-4550 using an *vitro* 3D cell proliferation assay (Fig. 1). The cell lines were selected based on their RAS mutational status, with the number of lines representing each onco-genotype weighted to reflect the prevalence of those mutations in human cancer genomes^18^.

**Figure 1.**
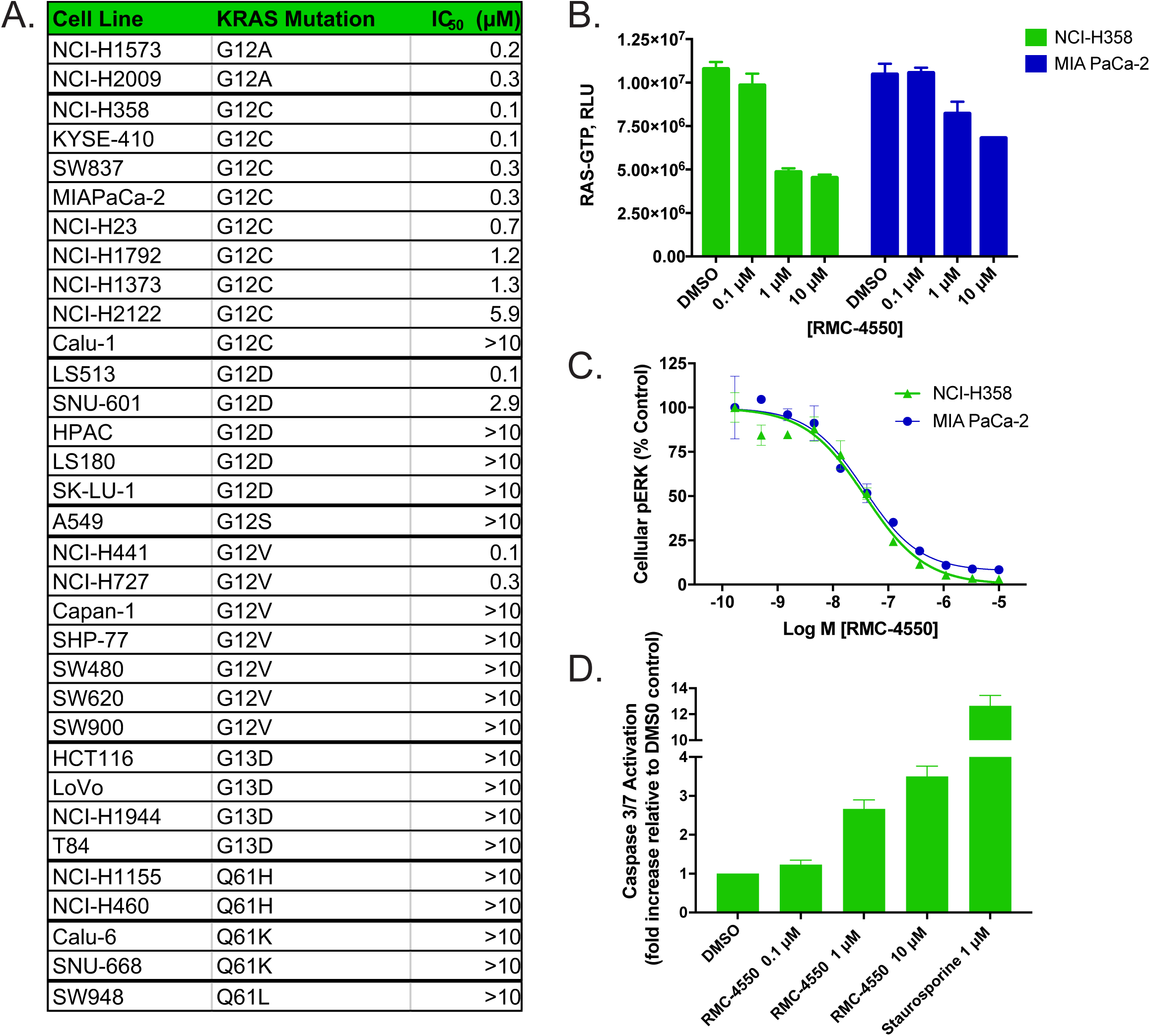
SHP2 Inhibition suppresses growth and RAS/MAPK signaling in a range of cancer cell lines driven by KRAS^G12^ mutations. A) Potency (IC_50_) of RMC-4550 for inhibition of proliferation of thirty-three KRAS mutant cancer cell lines grown in 3D culture. One day after seeding cells were treated with RMC-4550 and cell viability measured on Day 8 using CTG. IC_50_ values for growth inhibition were calculated as described in the methods. **B)** and **C)** Representative KRAS^G12C^ lines, NCI-H358 and MIA PaCa-2, were grown in 2D culture and incubated with increasing concentrations of RMC-4450 for one hour. Cellular lysates were prepared and levels of RAS-GTP **(B)** and pERK **(C)** determined. RMC-4550 produced a concentration-dependent reduction in both cellular RAS-GTP and p-ERK levels. Geometric mean IC_50_ values for reduction in pERK in H358 and MIAPaCa2 cells were 46 nM and 53 nM respectively (n = 2 observations, in duplicate). **D)** NCI-H358 cells were grown on ULA plates as spheroids. After 5 days in culture, spheroids were treated with RMC-4550 or staurosporine, as a positive control, and assayed for caspase 3/7 activity after 20 hours. RMC-4550 produced a concentration-dependent increase in caspase 3/7 activity.

Strikingly, cancer cell lines bearing missense mutations in KRAS at G12, but neither G13 nor Q61, exhibited sensitivity to RMC-4550 (IC_50_ < 2 μM, Fig. 1A). The observation that cancer cell lines bearing the G12C mutation were preferentially sensitive to SHP2 inhibition (hypergeometric p-value = 0.0043) is consistent with prior work demonstrating that the sensitivity of KRAS^G12C^ cancer cell lines to a selective KRAS^G12C^ inhibitor can be modulated by upstream signals that regulate RAS activity^9-11^. The proliferation of two G12C cell lines (NCI-H2122 and Calu-1) was insensitive to SHP2 inhibition. In Calu-1 cells, RMC-4550 potently suppressed pERK consistent with inhibition of pathway activation (IC_50_ = 7 nM, Fig. S3). The effect of RMC-4550 on pERK in NCI-H2122 cells was not tested. The differential sensitivity to growth inhibition across G12C cell lines may reflect intrinsic variability in the dependence of each cell line on signaling via the RAS/MAPK pathway. In addition, G12A mutant cell lines were sensitive to SHP2 inhibition, while cell lines bearing G12V and G12D mutations exhibited a range of responses. These data suggest that cancers bearing additional oncogenic KRAS variants may be sensitive to modulation by upstream factors^19,20^.

In NCI-H358 (lung, KRAS^G12C/+^) and MIA PaCa-2 (pancreas, KRAS^G12C/G12C^) cell lines, RMC-4550 not only inhibited cell proliferation, but strongly suppressed RAS-GTP and pERK levels (Fig. 1B,C, S4). These results support a model in which SHP2 functions upstream of certain nucleotide-cycling forms of oncogenic KRAS to regulate its activation by promoting or sustaining GTP-loading of RAS.

To characterize further the cellular effects of RMC-4550, we examined a marker of apoptosis. In NCI-H358 cells, treatment of spheroid cultures with RMC-4550 led to caspase 3/7 activation, indicative of a pro-apoptotic effect (Fig. 1D).

Collectively, these data demonstrate that certain oncogenic G12 variants of KRAS remain dependent on SHP2-mediated GTP-loading to promote downstream signaling and cell proliferation. Therefore, we hypothesized that additional oncogenic mutations that drive RAS/MAPK signaling in a manner that is dependent upon RAS nucleotide exchange may also confer sensitivity to SHP2 inhibition.

### Loss of the Tumor Suppressor NF1 Confers Sensitivity to SHP2 Inhibition

NF1 acts as a tumor suppressor via the RAS GAP activity of its GAP-related domain (GRD). Accordingly, loss of NF1 function has been shown to increase RAS-GTP levels, permit hyperactive RAS/MAPK signaling, and contribute to a variety of human cancers^5,6,21^. Because the increase in RAS-GTP levels is due to loss of RAS GAP function^22^ and wild-type RAS retains some level of NF1-independent GTPase activity that could attenuate RAS-GTP signaling^23^, we hypothesized that inhibition of RAS-GTP loading would offset the loss of RAS GAP activity and inhibit RAS-mediated downstream oncogenic signaling. Therefore we tested whether NCI-H1838, an NF1^LOF^ lung adenocarcinoma cell line (Fig. S5A), was sensitive to SHP2 inhibition.

Consistent with our hypothesis, treatment of H1838 cells with RMC-4550 led to downregulation of RAS-GTP levels, suppression of pERK, and inhibition of cell proliferation (Fig. 2A-C). In SK-MEL-113, an NF1^LOF^ melanoma line, we observed antiproliferative activity of another allosteric SHP2 inhibitor (Fig. S5B,C). These data indicate that loss of the tumor suppressor NF1 is a second class of downstream oncogenic mutation that can be targeted through inhibition of RAS-GTP loading.

**Figure 2.**
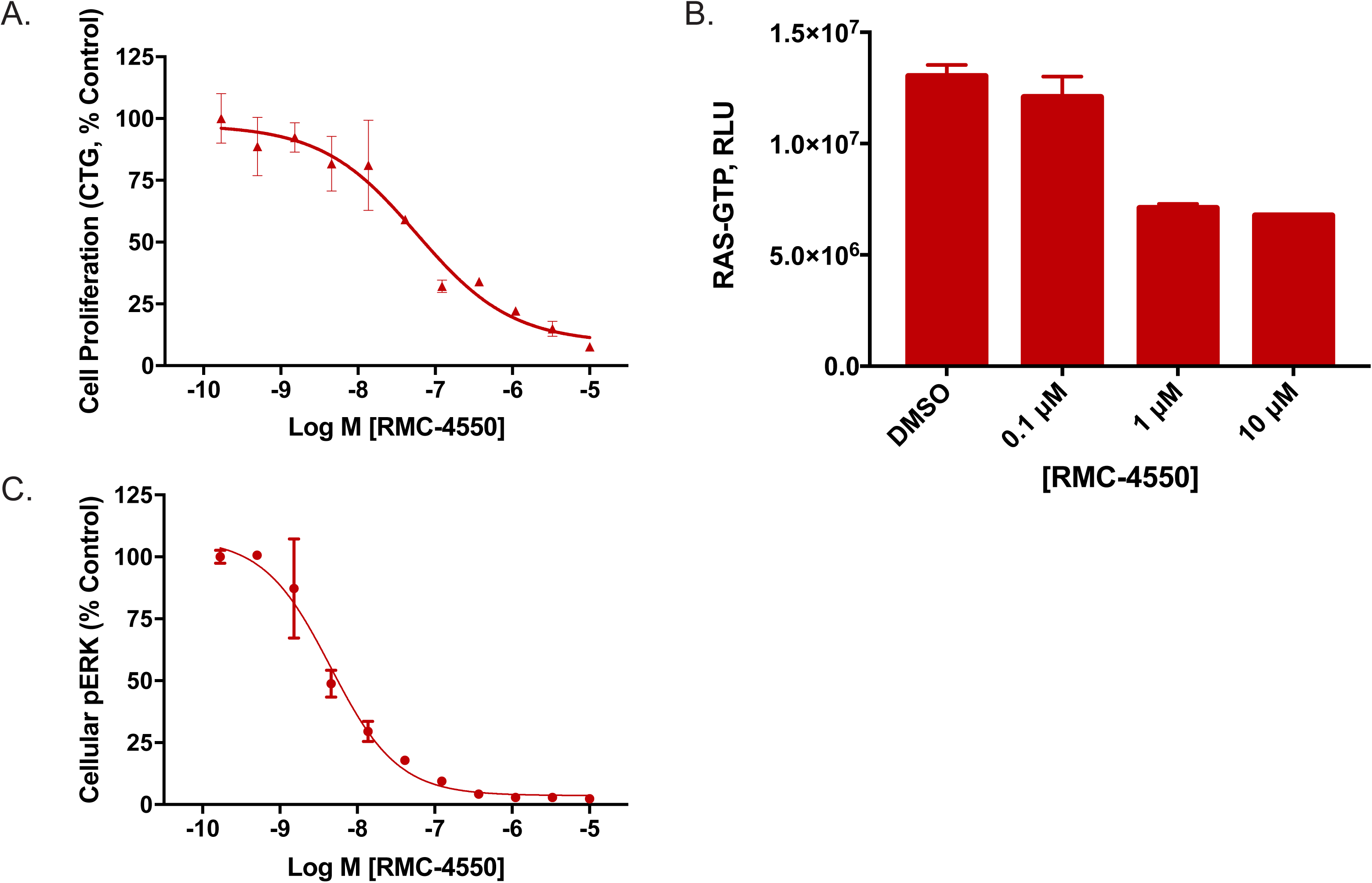
SHP2 inhibition suppresses growth and RAS/MAPK signaling in a cancer cell line driven by NF1^L0F^ mutation. A) Effect of RMC-4550 on proliferation of H1838 NF1^LOF^ cells in 3D culture. One day after seeding cells were treated with RMC-4550 and cell viability measured on Day 8 using CTG. Data are representative of 2 independent experiments each performed in duplicate. RMC-4550 exhibited a geometric mean IC_50_ value of 46 nM. B) and C) H1838 cells were grown in 2D culture and incubated with increasing concentrations of RMC-4450 for one hour. Cellular lysates were prepared and levels of RAS-GTP (B) and p-ERK (C) determined. RAS-GTP levels in H1838 cells were inhibited in a concentration-dependent manner by RMC-4550. The geometric mean IC_50_ value for reduction in pERK in H1838 cells was 4 nM (n = 3 observations, in duplicate).

### Class 3 BRAF Mutations Confer Sensitivity to SHP2 Inhibition

BRAF is a serine/threonine kinase that is frequently mutated in human cancers. Like oncogenic mutations in KRAS and loss of NF1, BRAF mutations drive cancer through hyperactivation of pERK signaling. There are at least three classes of oncogenic BRAF mutations that each differ in the mechanism by which they drive pERK signaling. Class 1 mutations occur at V600 and result in constitutively active monomers of BRAF that decouple BRAF from RAS-GTP activity^24^. Class 2 mutations preserve BRAF’s dependence on dimerization, but also decouple dimer-dependent signaling from RAS-GTP levels^25^. Therefore, we expected that cancer cell lines driven by either class 1 or class 2 mutations in BRAF would be insensitive to SHP2 inhibition. In contrast, the recently-described class 3 mutations of BRAF maintain RAF dimer and RAS-GTP dependence to drive pERK signaling^13^. In light of their dependence on RAS-GTP, we predicted that class 3 mutations of BRAF would be responsive to SHP2 inhibition of RAS-GTP levels. To test these predictions, we screened a representative panel of cell lines bearing oncogenic BRAF mutations across these three classes for sensitivity to SHP2 inhibition.

First, we confirmed that class 1 and 2 BRAF mutations were refractory to SHP2 inhibition (Fig. 3A-C). Consistent with the mechanistic framework described above, we observed that RMC-4550 did not inhibit pERK levels or proliferation in BRAF^V600E^ mutant A-375 cells. Similar results were observed in the NCI-H1755 cell line carrying a class 2 BRAF mutation (G469A), which exhibits RAS-independent homodimer formation and signaling^25^. Notably, RMC-4550 did not inhibit RAS-GTP levels in these cell lines. Class 1 and class 2 BRAF mutant oncoproteins function downstream of RAS but drive strong, ERK-dependent negative feedback that, through mechanisms upstream of RAS, leads to suppression of RAS-GTP^26^. Any residual activation of RAS that occurs in the presence of such negative feedback may either be insensitive to SHP2 inhibition, for example if suppression has occurred via direct inhibition of SOS1^27,28^, or below the level of detection for our RAS-GTP assay.

**Figure 3.**
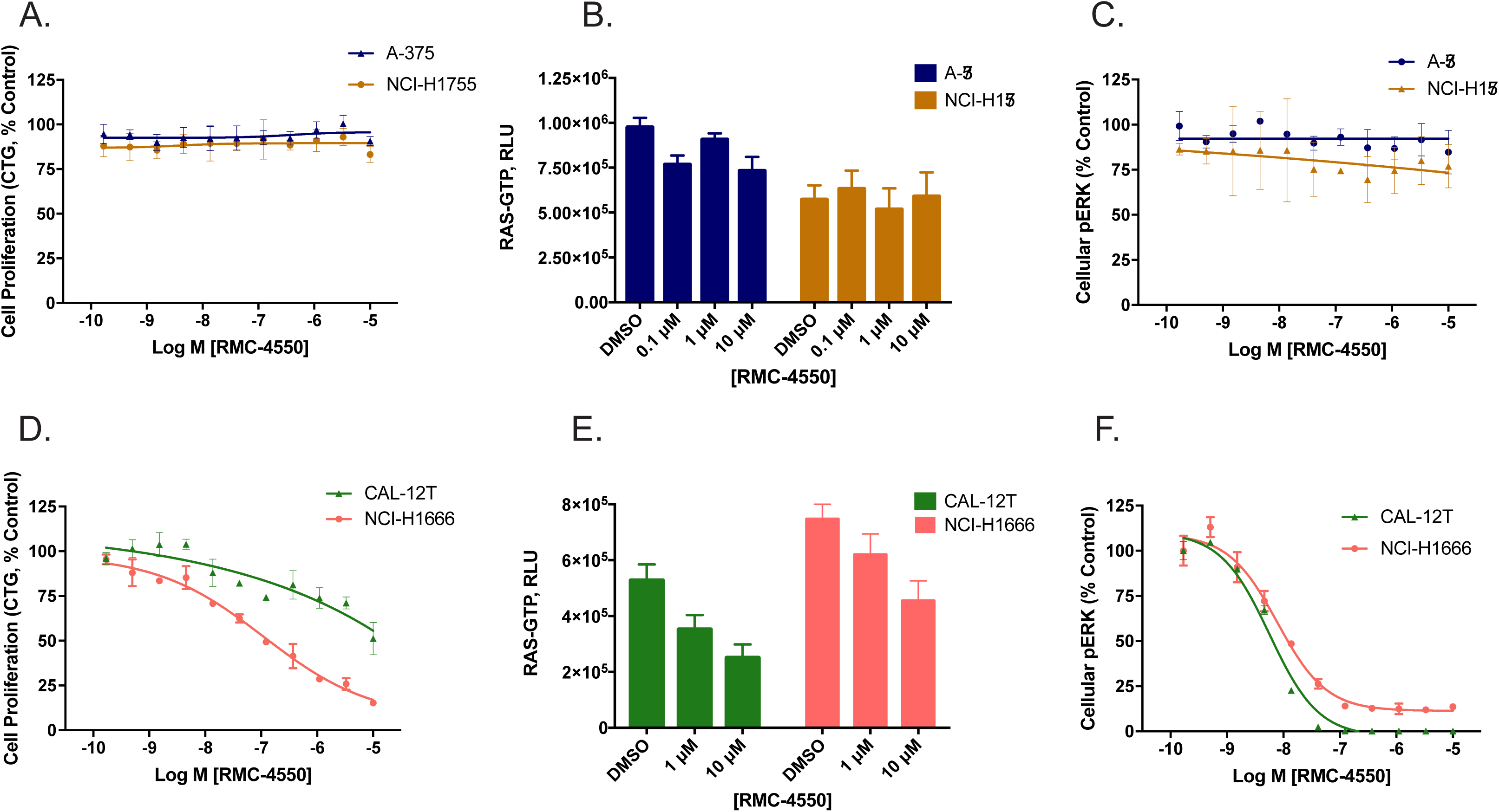
SHP2 inhibition suppresses growth and RAS/MAPK signaling in cancer cell lines with Class 3 BRAF mutations. A) Effect of RMC-4550 on proliferation of class 1 (A-375, BRAF^V600E^) and class 2 (NCI-H1755, BRAF^G469A^) BRAF mutant cell lines in 3D culture. Three days after seeding, cells were treated with RMC-4550 and cell viability measured on Day 8 using CTG. Data are representative of 2 independent experiments each performed in duplicate. RMC-4550 had no effect on proliferation up to the maximum test concentration of 10 μM. **B)** and **C)** A-375 and NCI-H1755 cells were grown in 2D culture and incubated with increasing concentrations of RMC-4450 for one hour. Cellular lysates were prepared and levels of RAS-GTP **(B)** and p-ERK **(C)** determined. RMC-4550 had no effect on cellular RAS-GTP or pERK levels in either cell line up to the maximal test concentration of 10 μM. Data are representative of 3 independent experiments each performed in duplicate. **D)** Effect of RMC-4550 on proliferation of two class 3 BRAF mutant cell lines (CAL-12T, BRAF^G466V/+^; NCI-H1666, BRAF^G466V/+^) cells in 3D culture. Three days after seeding, cells were treated with RMC-4550 and cell viability measured on Day 8 using CTG. Data are representative of 2 independent experiments each performed in duplicate. RMC-4550 exhibited geometric mean IC_50_ values of >10 μM and 236 nM respectively, for growth inhibition in CAL-12T and NCI-H1666 cells. **E)** and **F)** CAL-12T and NCI-H1666 cells were grown in 2D culture and incubated with increasing concentrations of RMC-4450 for one hour. Cellular lysates were prepared and levels of RAS-GTP **(E)** and p-ERK **(F)** determined. RAS-GTP levels in CAL-12T and NCI-H1666 cells were inhibited in a concentration-dependent manner by RMC-4550. RMC-4550 produced a concentration-dependent reduction in pERK levels in CAL-12T and NCI-H1666 cells with geometric mean IC_50_ values of 4 nM and 9 nM, respectively (n = 3 observations, in duplicate).

In contrast, in two cell lines carrying class 3 BRAF mutations, H1666 (BRAF^G466V/+^) and CAL-12T (BRAF^G466V/+^), treatment with RMC-4550 led to suppression of RAS-GTP levels, pERK levels, and, in the case of H1666, proliferation. These results are consistent with recent reports that class 3 BRAF mutations are bona fide cancer drivers that remain sensitive to modulation of upstream signaling and RAS-GTP levels^13^. Therefore, class 3 BRAF mutations are a third category of downstream oncogenic mutation that can be targeted through blockade of SHP2-mediated RAS-GTP loading.

### Inhibition of SHP2-dependent RAS-GTP loading can be rescued by constitutive activation of SOS1

In light of our findings that SHP2 inhibition can be used to target distinct classes of RAS/MAPK pathway oncoproteins that remain dependent upon RAS-GTP loading, we asked whether SHP2-dependent modulation of RAS-GTP was due to disruption of core RAS-regulatory processes. First, we mined data from the recently published Project DRIVE^29^, in which thousands of genes were systematically depleted across hundreds of cell lines to study genetic dependencies of molecularly defined cancer cell lines. One way to identify functional modules from high-throughput genetic knockdown experiments is to examine the phenotypic correlation of all possible gene pairs across the full dataset, as knockdown of members of a common functional module tends to yield similar patterns of response over many independent experiments. Taking a hypothesis-driven approach, we pulled data for twenty-three genes involved in RTK/RAS/MAPK signaling and calculated a correlation matrix (Fig. 4A). Two functional modules were readily apparent – the MAPK signal relay downstream of activated RAS and the RTK/convergent node module upstream of activated RAS. Consistent with the present observations, the most closely correlated knockdowns to PTPN11 (SHP2) are the GEF protein SOS1 (correlation coefficient =0.51) and the adaptor protein GRB2, which links RTKs to SOS1-mediated GTP-loading of RAS (correlation coefficient =0.40). In fact, SOS1 and GRB2 are the most closely related gene knockdowns to PTPN11 across all 7,837 genes in the Project DRIVE dataset^30^. Consistent with previous work, this analysis implies that SHP2 is an essential member of a core RAS-regulatory module containing SOS1 and GRB2. We therefore hypothesized that RMC-4550 downregulates RAS-GTP by disrupting the SHP2/SOS1/GRB2 module that is required for GTP-loading of RAS^31,32^.

**Figure 4.**
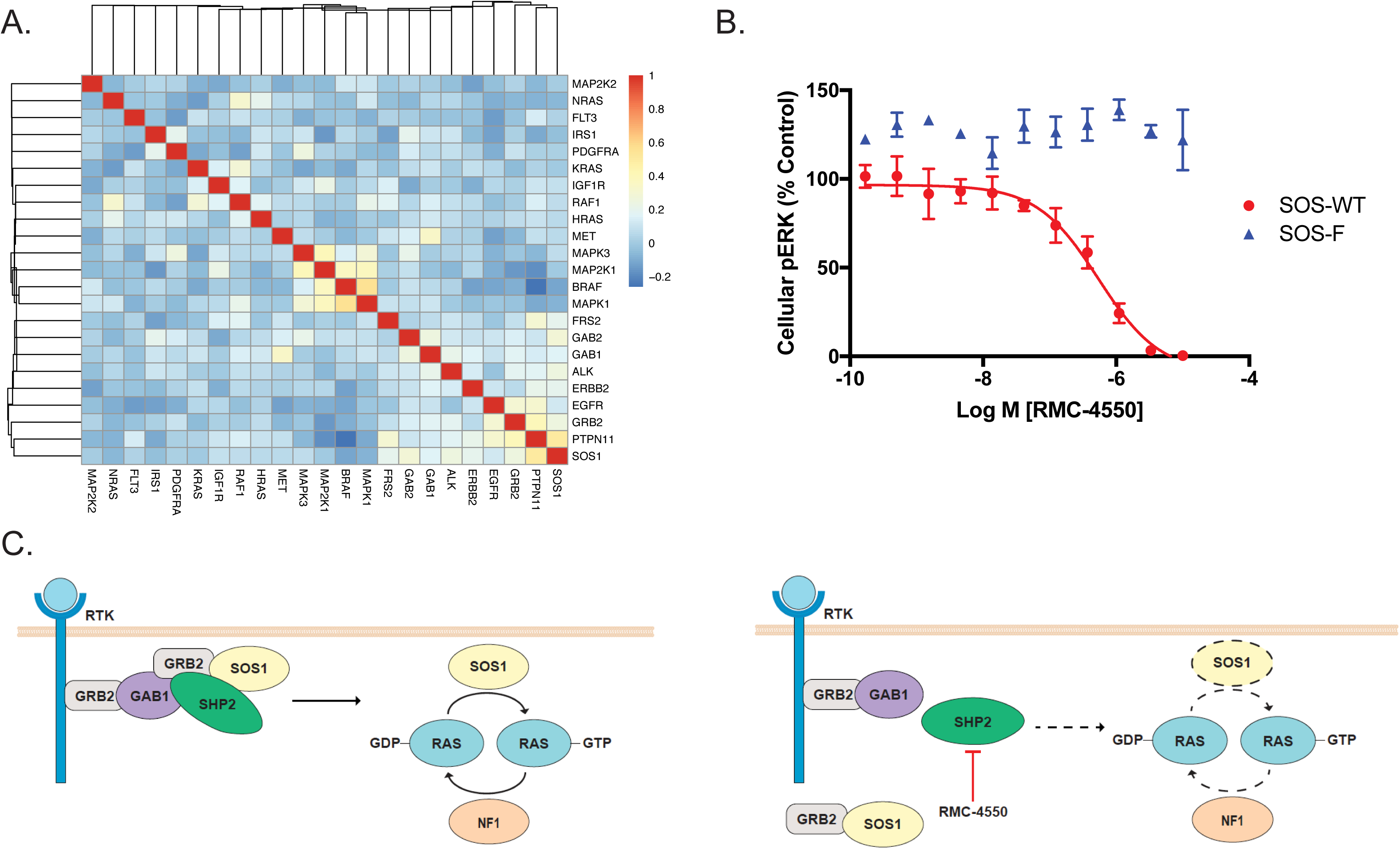
Mechanism of action of SHP2 inhibition is dependent on SOS1. A) Correlation analysis of the cellular effects of genetic knockdown of signaling molecules in the RTK/RAS/MAPK pathway in Project DRIVE. Knockdown of PTPN11 (SHP2) is most closely correlated with SOS1 (correlation coefficient 0.51) and GRB2 (correlation coefficient 0.4) suggesting these are all members of a core RAS-regulatory module **B)** Effect of RMC-4550 on cellular pERK in HEK293 expressing SOS-WT (wild type) or SOS-F, a SOS1 mutant that targets SOS protein constitutively to the plasma membrane. Expression of SOS-WT and SOS-F was induced with doxycycline for 24 hours. Cells were incubated for 1 hour with RMC-4550 and stimulated with EGF (50 ng/mL) for the final five minutes of drug treatment. Cellular lysates were prepared for determination of pERK. RMC-4550 produced a concentration-dependent reduction in pERK levels in cells expressing SOS-WT but not SOS-F, geometric mean IC_50_ = 1.0 μM (n = 3 observations, performed in duplicate). In SOS-F expressing cells, no inhibition was observed up to maximal test concentration of 10 μM. **C)** Cartoon representation of the function of SHP2, in conjunction with GRB2, as a transducer of signals from upstream RTKs to SOS1 for RAS activation.

To examine this hypothesis, we tested whether a dominant, constitutively-active mutant form of SOS1 could render cells insensitive to RMC-4550-mediated inhibition of RAS/MAPK signaling. Indeed, in HEK293 cells, inducible expression of SOS-F, a SOS1 mutant in which the C-terminal region containing a GRB2 binding site is replaced with the HRAS farnesylation motif that targets the protein constitutively to the plasma membrane^33^, rendered cells insensitive to either EGF stimulation (Fig. S6) or SHP2 inhibition of pERK (Fig. 4B). These data show that the effects of SHP2 inhibition can be bypassed by constitutive activation of SOS1 and that SOS1 therefore functions downstream of (or in parallel to) PTPN11/SHP2. The precise mechanistic role of SHP2 in MAPK signaling is still incompletely understood. One possible explanation for these findings is that allosteric inhibition of SHP2 interferes with the formation of a multi-protein complex at the RTK through disruption of productive binding of the SHP2 SH2 domains with adaptor proteins such as GAB1 (Fig. 4C). Phosphorylated tyrosines in the C-terminal tail of SHP2 are proposed to bind GRB2 and may modulate SOS1 plasma membrane localization via the well-established GRB2/SOS1 association^34,35^. We would expect potential disruption of this interaction by RMC-4550 to be bypassed by SOS-F, which is targeted to the membrane independently of GRB2 association. Alternatively, we cannot rule out that inhibition of SHP2 phosphatase activity drives decreased pERK levels through modulation of SH2 domain-binding sites or other regulatory domains in SHP2 target proteins.

### SHP2 inhibition by RMC-4550 treatment suppresses tumor growth and oncogenic signaling in models of KRAS^G12C^-driven cancer *vivo*

Building on our *vitro* Normalized valuefindings, we asked whether the sensitivity to SHP2 inhibition observed in a majority of KRAS^G12C^ mutant cell lines was recapitulated *vivo*. To this end, we examined the effect of increasing doses of RMC-4550 on tumor growth in NCI-H358 (KRAS^G12C/+^) and MIA PaCa-2 (KRAS^G12C/G12C^) xenograft tumors in immune-deficient mice. We observed dose-dependent efficacy following daily oral dosing of RMC-4550 in each of these models (Figure 5A, B) with tumor growth inhibition observed at all doses. All doses of RMC-4550 were well tolerated in both experiments (Fig. S7).

**Figure 5.**
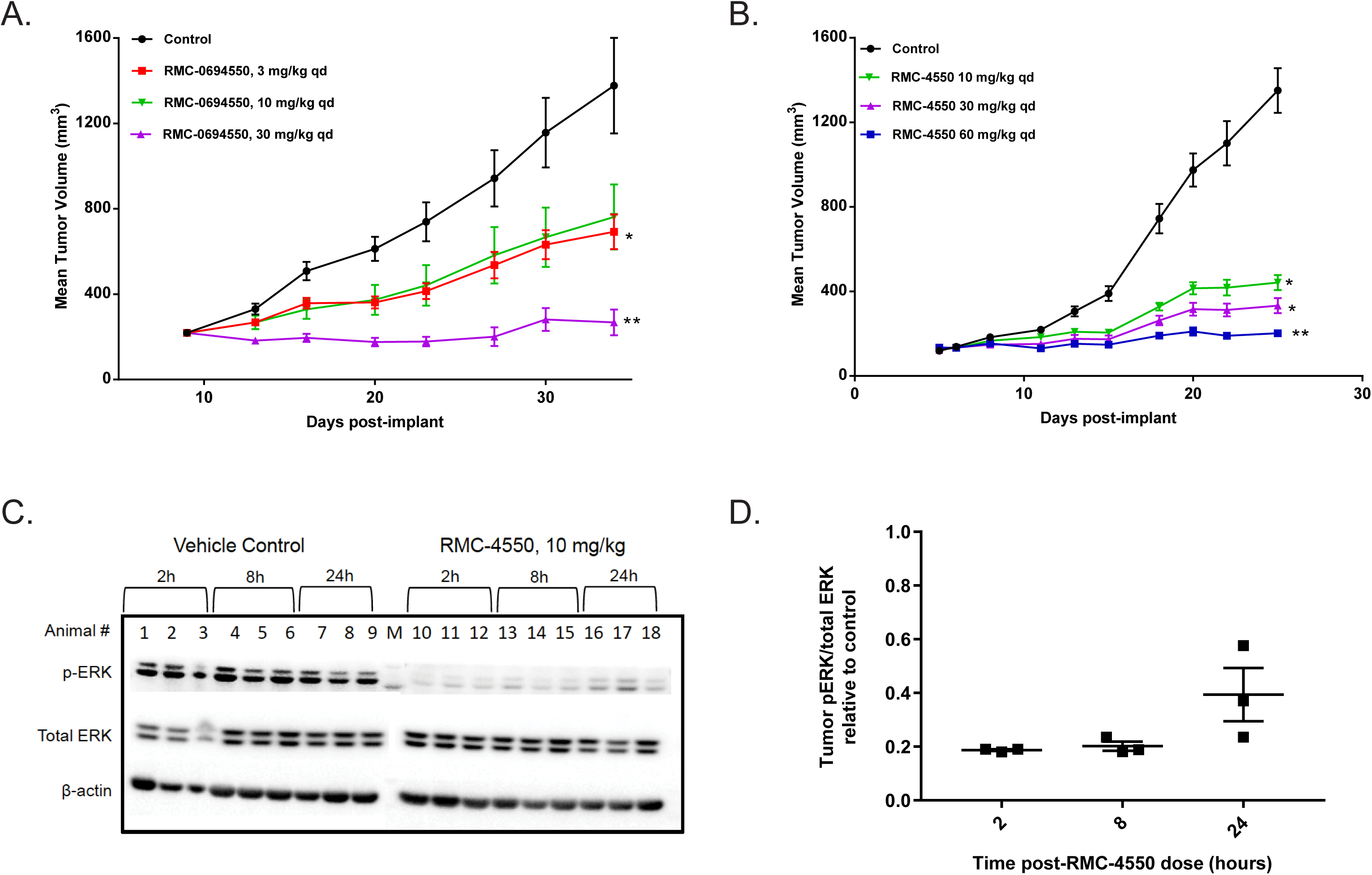
SHP2 inhibition suppresses tumor growth and RAS/MAPK signaling in xenograft models of KRASG12C-driven cancer. Daily oral administration of RMC-4550 produced a dose-dependent decrease in tumor volume in two xenograft tumor models. **A)** NCI-H358 (n = 10/group) showed tumor growth inhibition (TGI) at 3, 10, and 30 mg/kg (TGI = 59, 53, 96% respectively) and **B)** MIA PaCa-2 (n = 12/group) exhibited significant tumor growth inhibition at 10, 30 and 60 mg/kg (TGI = 74, 83, 93% respectively). * p < 0.05, ** p < 0.01, ordinary one way ANOVA with post-hoc Tukey’s test. Figures show mean +/- S.E.M. **C)** Representative Western blot showing relative suppression of pERK in NCI-H358 tumors over a period of 24 hours following a single dose of 10 mg/kg. Maximal suppression relative to controls was observed at 2 and 8 hours following the dose, with some recovery at 24 hours. **D)** Quantitative densitometric analysis of Western blot shown in C. pERK/total ERK ratio in treated tumors normalized to mean ratio in control tumors (set at 1) at each time point (n = 3/group). Figure shows mean +/- S.E.M.

The pharmacodynamic effects of RMC-4550 were explored in the NCI-H358 (KRAS^G12C/+^) model. We observed time-dependent suppression of pERK over a period of 24 hours following a single oral dose of RMC-4550 in these xenograft tumors (Fig. 5C, D). This pharmacodynamic effect is consistent with the dose-dependent efficacy observed upon repeated daily dosing.

## DISCUSSION

We have discovered a novel, potent and selective allosteric SHP2 inhibitor, RMC-4550, and used it to identify molecular markers of SHP2-dependence in tumors bearing mutations in the RAS/MAPK pathway. The observation that KRAS^G12C^, NF1^LOF^, and class 3 BRAF mutations each confer sensitivity to SHP2 inhibition in tumor cells establishes SHP2 inhibition as a novel and promising therapeutic strategy against tumors that harbor these oncogenic driver mutations.

In non-small cell lung cancer (NSCLC), these driver mutations are observed frequently: KRAS^G12C^, NF1^LOF^, and class 3 BRAF mutations collectively represent approximately 20-25% of all cases annually^4,36^. Importantly, patients whose cancers carry these mutations are dramatically underserved, as no targeted therapies have been approved for these molecular subtypes. The data presented here raise the exciting possibility that a SHP2 inhibitor may convert these mutations into clinically-actionable targets and improve clinical outcomes.

Our data show that SHP2 is a convergent signaling node downstream of multiple RTKs and an essential regulator of oncogenic RAS activation. Importantly, some tumors remain sensitive to SHP2 inhibition even when the oncogenic ‘driver’ mutation is apparently downstream of SHP2 in the canonical pathway, as for class 3 BRAF mutations. The known association of SHP2 with SOS1 and GRB2 provides a mechanistic context for SHP2’s precise role in the regulation of RAS-GTP levels, and suggests that allosteric inhibitors of SHP2 disrupt this functional module that is essential for RAS activation. Further mechanistic studies are warranted to dissect how SHP2 inhibition interferes with RAS-GTP loading in the cancers with KRAS^G12C^, NF1^LOF^, and class 3 BRAF mutations. A previous study reported that RAS mutants as a class functioned independently of SHP2 and concluded that cells carrying mutations of downstream proteins, including RAS and RAF, broadly were insensitive to SHP2 inhibition^17^. There may be several reasons for this discrepancy, including differences in both the specific oncogenic mutants studied and methodologies. Nonetheless, the present findings demonstrate both functionally and mechanistically that SHP2 is a relevant target for modulating select downstream oncoproteins including specific variants of RAS and BRAF.

The dependence of KRAS^G12C^, NF1^LOF^, and class 3 BRAF mutations on SHP2-mediated upstream signals suggests that certain mutant forms of RAS pathway oncogenic drivers amplify, rather than bypass, the physiologic signals and homeostatic mechanisms regulating RAS-GTP and pathway output. This model contrasts with a common assumption that RAS oncogenes are locked in the GTP-bound “on” state constitutively to drive signaling and cancer, and instead is consistent with an emerging framework in which certain oncogenic mutations are semi-autonomous, rather than fully-autonomous, drivers of cancer. Accordingly, our findings illuminate previously unrecognized oncology indications for SHP2 inhibition based on patient selection beyond cancers with RTK alteration. More broadly, our study highlights the power of developing selective and potent pharmacologic probes to uncover occult features of oncogenic RAS signaling and unanticipated therapeutic opportunities.

## MATERIALS AND METHODS

### Cell Lines and Reagents

Cell lines were purchased from the American Type Culture Collection (ATCC) and maintained at 37°C in a humidified incubator at 5% CO2. Cells were grown in appropriate media as recommended by ATCC, and supplemented with 10% fetal bovine serum (Cellgro) and 1% penicillin/streptomycin (Gibco) unless directed otherwise.

### RAS-mutant Cancer Cell Panel Screen

Cell lines were maintained in growth medium supplemented with 10% fetal bovine serum and 1% penicillin/streptomycin at 37°C, 5% CO_2_. For each line, two 96-well plates were seeded with 600 cells/well suspended in 0.65% methylcellulose. Plates were incubated at 37°C, 5% CO_2_. The following day, one of the plates for each line was used to obtain initial (T=0) readings using the CellTiter-Glo (CTG) assay kit (Promega), following the manufacturer’s instructions. Luminescence was read in an EnVision Multilabel Plate Reader (Perkin Elmer). Serial 3-fold dilutions of RMC-4550 were prepared in growth medium supplemented with 10% fetal bovine serum and 1% penicillin/streptomycin (final DMSO concentration = 0.1%) and added to each well. Cells were incubated for seven days, and the number of viable cells was quantitated using the CTG assay, as above. Day 7 (T=7) values for a given RMC-4550 concentration *X* were individually normalized against T=0 values according to the formula: Normalized value = ((S_7_[*x*]-S_0_)/(S_7_[DMSO]-S_0_))*100, where S_7_ is the CTG signal at T=7, and S_0_ is the CTG signal at T=0. Normalized data was imported into Prism 7 (GraphPad) software for plotting and calculation of IC_50_ values using four-parameter concentration-response model.

### Analysis of ERK1/2 Phosphorylation

ERK1/2 phosphorylation at Thr202/Tyr204 was assayed using the AlphaLISA SureFire Ultra HV pERK Assay Kit (Perkin Elmer). 24 hours prior to the assay, 20,000 cells per well were plated in clear 96-well plates, in biotin-free media supplemented with 10% fetal bovine serum and 1% penicillin/streptomycin, and incubated overnight at 37°C in 5% CO_2_. One hour prior to treatment with RMC-4550, media was replaced with biotin-free media supplemented with 0.02% bovine serum albumin and 1% penicillin/streptomycin. Cells were incubated at 37°C in 5% CO_2_ for one hour. Cells were then exposed to serial 3-fold dilutions of RMC-4550 diluted in biotin-free media supplemented with 0.02% bovine serum albumin and 1% penicillin/streptomycin (final DMSO concentration equivalent to 0.1%). Cells were incubated in the presence of RMC-4550 for one hour at 37°C in 5% CO_2_. Cellular lysates were prepared and pERK levels determined according to the manufacturer’s protocol. Samples were read using an EnVision Multilabel Plate Reader (Perkin Elmer) using standard AlphaLISA settings. Assay data was plotted and EC_50_ values were determined using four-parameter concentration-response model in GraphPad Prism 7. Data in figures are mean +/- standard deviation of duplicate values from representative experiments.

### Analysis of RAS-GTP Levels

Levels of activated RAS-GTPase were determined using the Ras GTPase ELISA Kit (Abcam), according to the manufacturer’s instructions. Briefly, 1x10^6^ cells were seeded in media supplemented with 10% fetal bovine serum and 1% penicillin/streptomycin in 10 cm tissue culture dishes, and incubated for 48 hours at 37°C in 5% CO_2_. Following this incubation period, cells were treated with RMC-4550 diluted in media supplemented with 0.02% bovine serum albumin and 1% penicillin/streptomycin to a final DMSO concentration equivalent to 0.1%. Cells were incubated in the presence of RMC-4550 for one hour at 37°C in 5% CO_2_. Following this exposure period, media was removed and cells were washed once in ice-cold PBS. Cells were immediately lysed in ice-cold 1x Lysis buffer (25 mM Tris-HCl, pH 7.2, 150 mM NaCl, 5 mM MgCl2, 1% NP-40, 5% glycerol)(Thermo) supplemented with Halt Protease and Phosphatase Inhibitor (EDTA-free)(Thermo), and lysates were cleared by centrifugation at 13,000 rpm for 15 minutes at 4°C. Protein quantification was performed with the BCA Protein Assay Kit (Pierce), and lysates were diluted to a final concentration of 1 mg/mL. Cell lysates were prepared and levels of RAS-GTP determined using a plate-based ELISA as described in manufacturer’s protocol. Data in figures are mean +/- standard deviation of duplicate values from representative experiments.

### Spheroid Formation and Proliferation

2500 cells/well were seeded in round bottom ultra-low attachment 96-well plates (Corning) in growth media supplemented with 10% fetal bovine serum and 1% penicillin/streptomycin, and allowed to form spheroids for 72 hours at 37°C in 5% CO_2_. Spheroid formation was confirmed visually, and spheroids were treated in duplicate with serial 3-fold dilutions of RMC-4550 in complete growth media (final DMSO concentration = 0.1%). Following drug exposure for five days, cell viability in spheroids was determined using the CellTiter-Glo assay kit (Promega), following the manufacturer’s instructions. Luminescence was read in a SpectraMax M5 Plate Reader (Molecular Devices). Assay data was normalized to DMSO values, and EC_50_ values were determined using a four-parameter concentration-response model in GraphPad Prism 7. Data in figures are mean +/- standard deviation of duplicate values from representative experiments.

### Activated Caspase 3/7 Assay in Spheroids

H358 cells were grown into spheroids by seeding 5,000 cells/well in round bottom ultra-low attachment 96-well plates (Corning) in RPMI media supplemented with 10% fetal bovine serum and 1% penicillin/streptomycin and incubated at 37°C in 5% CO_2_ for five days to allow for spheroid formation. Spheroid formation was confirmed visually. Spheroids were treated in triplicate with RMC-4550, staurosporine (Sigma), or DMSO (Sigma) (0.1% final), diluted in RPMI media supplemented with 10% fetal bovine serum and 1% penicillin/streptomycin, and incubated at 37°C in 5% CO_2_ for 20 hours. Caspase 3/7 activity was measured using the Caspase-Glo 3/7 Assay System (Promega), following the manufacturer’s instructions. Luminescence was read in an EnVision Multilabel Plate Reader (Perkin Elmer).

### Gene Depletion Sensitivity Correlation Analysis

Gene signatures were downloaded from the Project Drive data portal https://oncologynibr.shinyapps.io/drive) and assembled into a matrix. Correlation analysis was conducted in R and heatmaps generated with the pheatmap package.

### SOS-WT and SOS-F Expression Constructs

N-terminally HA-tagged SOS-WT and SOS-F constructs were synthesized (Atum) and subcloned into the pcDNA5/FRT/TO vector (ThermoFisher) using the following primers: SOS1-HA-For 5’-ACAGGTAAGCTTATGTACCCATACGATGTTCCAGATTAC-3’, SOS1-HA-REV 5’-AGACTAGCGGCCGCTCAGGAAGAATGGGCATTCTCCAA-3’, and SOS-F-HA-REV 5’-GATCGAGCGGCCGCTCAGGAGAGCACACACTTGCAG-3’. SOS-WT and SOS-F plasmids were co-transfected with the pOG44 Flp-recombinase expression vector (ThermoFisher) into the HEK Flp-In T-Rex 293 cell line according to the manufacturer’s protocol. Transfected cells were selected in drug media (200 μg/mL hygromycin B, 15 μg/mL blastidicin) and expression of SOS constructs was verified by western blot (SOS-1: Cell Signaling Technologies æ 5890; HA: Sigma 11867423001).

### pERK Analysis of HEK-293 SOS-WT and SOS-F

30,000 HEK-293 cells per well were plated in 96-well plates in Biotin-free RPMI (Hyclone) supplemented with 0.1% fetal bovine serum, 0.02% bovine serum albumin and 1% penicillin/streptomycin. Expression of SOS1 constructs was induced by the addition 0.1 μg/mL doxycycline (Sigma) for 24 hours. Cells were treated with serial 3-fold dilutions of RMC-4550 diluted in biotin-free media supplemented with 0.02% bovine serum albumin and 1% penicillin/streptomycin (final DMSO concentration equivalent to 0.1%) for one hour. For the final 5 minutes of drug treatment, cells were stimulated with 50 ng/mL EGF (Sigma), lysed and subjected to ERK1/2 phosphorylation analysis as described above.

### Purification of SHP2 FL and SHP2 cat Proteins

Constructs for the SHP2FL (Met1‐ Arg593) and SHP2Cat (Phe247-Arg529) were generated by cloning the *PTPN11*gene into a modified pET15 plasmid (GeneArt) containing an N-terminal 6x Histidine tag. The constructs were transformed into BL21 (DE3) cells and grown at 37 °C in Luria Bertani broth (LB) containing 50 μg/ml ampicillin. At an OD_600_ 0.8 cells were transferred at 18 °C and expression was induced by the addition of 0.5 mM IPTG. Cells were collected by centrifugation after overnight growth at 18 °C.

Cell pellets were resuspended in lysis buffer containing 25 mM Tris-HCl pH 8.0, 300 mM NaCl, 5 mM Imidazole, 1 mM TCEP, 1 μg/ml DNase and Complete EDTA-free protease inhibitors. The solution was incubated at 4 °C with stirring for 30 minutes. After the cell pellet was completely suspended the cells were sonicated on ice and centrifuged at 40,000 g for 45 min. The supernatant was loaded under gravity to a 5ml Ni NTA column equilibrated with 25 mM Tris-HCl pH 8.0, 300 mM NaCl, 5 mM Imidazole and 1 mM TCEP. The column was washed with the above buffer until the absorbance at 280nm (A_280_) was close to zero. Protein was eluted with buffer containing 25 mM Tris-HCl pH 8.0, 300 mM NaCl, 300 mM Imidazole and 1 mM TCEP. Fractions were collected and their protein content was determined by A280. Fractions containing either SHP2FL or SHP2Cat were pooled and concentrated with Amicon YM30 (SHP2FL) or YM10 (SHP2Cat) centrifugal concentrator. Concentrated protein was loaded to a Superdex 200 Increase 10/300 GL column equilibrated with 25 mM Tris-HCl pH 8.0, 150 mM NaCl and 3mM TCEP.

### SHP2 inhibition assay

Full length SHP2 was activated using a bisphosphorylated peptide derived from GAB1 (sequence H_2_N-VE(pY)LDLDLD(PEG8)RVD(pY)VVVDQQ-amide). The catalytic activity of SHP2 was monitored using the fluorogenic small molecule substrate DiFMUP (ThermoFisher) in 96-well, black polystyrene plates (Corning). The final reaction volume was 100 μl, and the final assay conditions were 55 mM HEPES pH 7.2, 100 mM NaCl, 0.5 mM EDTA, 1 mM DTT, 0.001% Brij35, 0.002% BSA, 0.1% DMSO, 20 μM DiFMUP, 0.2 nM enzyme, 500 nM activating peptide, and 10 μM to 1.9 pM inhibitor. Inhibitors were dissolved and serial 3-fold dilutions were prepared in DMSO. DMSO solutions were then diluted 1:100 in 50 mM HEPES pH 7.2 with 0.02% BSA. Enzyme was mixed with activating peptide in 2X buffer 10-60 minutes before starting the experiment. Diluted compound (10 μL) was mixed with activated enzyme (50 μl) and incubated for 30 minutes at room temperature. A 50 μM aqueous DifMUP solution (40 μl) was added and the plate was read in kinetic mode on a SpectraMax M5 plate reader (Molecular Devices) for 6 minutes using excitation and emission wavelengths of 340 nm and 450 nm. Plots of fluorescence units vs. time were fit with linear regression to determine initial velocity. Plots of initial velocity vs. inhibitor concentration were fit using a four-parameter concentration-response model in GraphPad Prism 7. For display purposes the concentration response curves were normalized using uninhibited and maximally inhibited controls on the experimental plates. The inhibition assay for the isolated catalytic domain of SHP2 was identical except that the activating peptide was omitted.

### Tumor Xenograft and *vivo* Pharmacodynamics

All animal research studies were conducted in accordance with institutional guidelines as described in approved IACUC protocols. Female (6-8 week old) athymic nude mice were implanted with NCI-H358 (Balb/c strain background) or MIA PaCa-2 (NCR nude strain background) tumor cells subcutaneously in the flank. Once tumors reached an average size of ∼200 mm^3^, administration of RMC-4550 (3 to 60 mg/kg PO) or vehicle (Captisol/ 50 mM acetate buffer pH4.6 (10%/90%, w/v)) was initiated. RMC-4550 was dosed daily via oral gavage at specified doses and was well tolerated (Figure S5). For efficacy studies, animals were euthanized when a mean control tumor volume of ∼1500 mm^3^ was reached at day 25 (NCI-H358) or day 22 (MIA PaCa-2). For pharmacodynamics studies, animals received a single dose of either RMC-4550 or vehicle, followed by euthanasia at a specified time (2, 8 or 24 hours post-dose). At this time tumors were collected, minced and snap frozen immediately for protein isolation for Western blot analysis.

### Phosphatase Selectivity Panel

A panel of twelve protein tyrosine phosphatases was expressed with 6-his tags in E. coli and purified using standard Ni^2+^ affinity purification procedures. Two protein serine/threonine phosphatases (PP1 and PP2A) were also used and obtained from SignalChem. The inhibition assays for each phosphatase were similar to the conditions described for the SHP2 full length and catalytic enzymes, except that the assay was performed in 384 well black polystyrene plates (Corning) with a total well volume of 50 μl, and fluorescence was read as an endpoint after a five minute incubation at room temperature. The following additional enzyme-specific changes were made. The enzyme concentration was 0.3 nM (PTP1b, HePTP, Lyp, CD45, and PTPRB), 125 nM (LMPTPA and PRL1), 15 nM (LMPTPB), 5 nM (DUSP22 and STEP), 1.6 nM (PP1), 0.5 nM (PP2A), 0.25 nM (SHP1 catalytic domain), and 0.2 nM (SHP1 full length). The reaction time for PP1 and DUSP22 was 20 minutes, and 90 minutes for PRL1. For PP1 and PP2A the EDTA in the assay buffer was replaced with 5 mM MnCl_2_. The assay for full length SHP1 was similar to the assay for full length SHP2, except that the enzyme was activated with a synthetic activating peptide with sequence H_2_N-LN(pY)AQLWHA(PEG8)LTI(pY)ATIRRF-amide at 250 nM.

### Kinase selectivity panel

Kinase selectivity was determined using the ScanMax panel, DiscoverRX (42501 Albrae Street, Fremont, CA 94538, United States).

### Cellular targets selectivity panel

Off-target profile was determined using Eurofins SafetyScreen44 panel, Eurofins (Eurofins Cerep, Le Bois l’Evêque, B.P. 30001, 86 600 Celle l’Evescault, France).

## ACKNOWLEDGMENTS

Revolution Medicines thanks Steve Kelsey for scientific guidance and critical review of the manuscript, Kevan Shokat for scientific guidance, Nidhi Tibrewal for development of the phosphatase selectivity panel, and Jeff Jasper for scientific input.

